# Y-chromosome structural diversity in the bonobo and chimpanzee lineages

**DOI:** 10.1101/029702

**Authors:** Matthew T. Oetjens, Feichen Shen, Sarah B. Emery, Zhengting Zou, Jeffrey M. Kidd

## Abstract

The male specific regions of primate Y-chromosomes (MSY) are enriched for multi-copy genes highly expressed in the testis. These genes are located in large repetitive sequences arranged as palindromes, inverted-, and tandem-repeats termed amplicons. In humans, these genes have critical roles in male fertility and are essential for the production of sperm. The structure of human and chimpanzee amplicon sequences show remarkable difference relative to the remainder of the genome, a difference that may be the result of intense selective pressure on male fertility. Four populations of common chimpanzees have undergone extended periods of isolation and appear to be in the early process of speciation. A recent study found amplicons enriched for testis-expressed genes on the primate X-chromosome the target of hard selective sweeps, and male-fertility genes on the Y-chromosome may also be the targets of selection. However, little is understood about Y-chromosome amplicon diversity within and across chimpanzee populations. Here, we analyze 9 common chimpanzee (representing three subspecies: *Pan troglodytes schweinfurthii, Pan troglodytes ellioti*, and *Pan troglodytes verus*) and two bonobo (*Pan paniscus*) male whole-genome sequences to assess Y ampliconic copy-number diversity across the Pan genus. We observe that the copy-number of Y chromosome amplicons is variable amongst chimpanzees and bonobos, and identify several lineage-specific patterns, including variable copy-number of azoospermia candidates *RBMY* and *DAZ*. We detect recurrent switchpoints of copy-number change along the ampliconic tracts across chimpanzee populations, which may be the result of localized genome instability or selective forces.

## Introduction

In mammals, males paternally inherit the Y chromosome, which contains the genetic content for canalizing embryos toward masculine development. As would be expected given the male-specific inheritance, many Y-linked genes are spermatogenesis factors with ubiquitous expression in the testis (Lahn, et al. 2001). These spermatogenesis genes are multi-copy and found in long stretches of repetitive DNA termed amplicons (Hughes, et al. 2010; Skaletsky, et al. 2003). Interchromosomal duplications of amplicons on the Y resulted in the formation of amplicon families, which share a high degree of identity among members. Amplicons are found on the Y as tandem repeats, inverted repeats, and palindromes. These formations can have mutational consequences for the genes they harbor. Palindromes and inverted repeats introduce large hairpin formations that physically align homologous sequence. In this arrangement, gene conversion between amplicon arms can occur, which may reverse deleterious mutations found within the multi-copy genes (Rozen, et al. 2003). The majority of the Y does not undergo recombination with the X. Therefore, in the absence of a mechanism for rescue, the transmission and persistence of Y-linked deleterious mutations in a population would be inevitable (Bachtrog 2013). The amplicons are thought have to evolved in response to strong selection of spermatogenesis traits in mammals as protection against deleterious mutation. On the other hand, tandem repeats introduce a susceptibility of amplification or deletion of the interstitial sequence through non-allelic homologous recombination (NAHR), which is a frequent cause of gain or loss of genes in the human genome (Gu, et al. 2008; Kidd, et al. 2010).

Amplicons undergo constant change in the human lineage with structural polymorphisms accruing at ~10,000 times the rate of nucleotide polymorphism (Repping, et al. 2006). Many of these structural polymorphisms are located in the amplicon regions termed AZF b and c (azoospermia factor regions b and c). Large deletions of the AZFc (1.6-3.5Mb) are acquired through NAHR and strongly associated with male infertility (Rozen, et al. 2012). The genes within the AZFc include specific copies of *DAZ* (Deleted in Azoospermia), *CDY* (Chromodomain Y), *BPY2* (Basic Protein Y, 2), and *PRY* (PTPN13-Like, Y-Linked). Each of these genes is found in a different amplicon family named after the colors used to denote them on initial Y-chromosome assemblies: *DAZ* is found in the red amplicon, *CDY* in yellow, *BPY2* in green, and *PRY* in blue (Hughes and Page 2015). The most penetrant AZFc deletion reported is rare (1/2,320) and is caused by a NAHR event between two blue arms (b2/b4) and increases the risk of azoospermia by 145 fold (Rozen, et al. 2012). The critical genes in the AZFb region are copies of *RBMY* (RNA-Binding Motif Protein Y) located in the teal amplicon and additional copies of *CDY*. While deletions within the AZFb appear to be more rare in human populations and contribute less to the overall genetic contribution of azoospermia, AZFb deletions have been reported in patients with spermategenic failure (Longepied, et al. 2010). These epidemiological studies suggest that the amplicons harbor genes essential for the progression of germ cells to haploid stages and that the copy number of these genes has functional consequences.

The sequencing of a Western Chimpanzee (*Pan troglodyte verus*) Y revealed gross structural differences with the human sequence, specifically in the amplicon regions. The chimpanzee amplicon region is richer in content, in terms of the number unique families and copy number of members. The combined length of the chimpanzee amplicon regions are 14.7 Mb compared to the human amplicon regions, which is approximately 10.2 Mb in sequence. Interestingly, some of the AZFc amplicons and genes were not equivalent in copy number in the two primate species, including: *BPY2* (human reference: 3, chimp reference: 2), *CDY* (human reference: 4, chimpanzee reference: 5), and *PRY* (human reference:2, chimpanzee reference: 0). In contrast, *DAZ* and *RBMY* were found at the same copy number in the chimpanzee and human reference assembly, 4 and 6, respectively.

The Western chimpanzee is one of four recognized subspecies, which based on a recent Y-chromosome inferred phylogeny diverged from a common ancestor 1.15 mya (Hallast, et al. 2016). The Central chimpanzee of Central Africa (*Pan troglodytes troglodytes*) shows genetic evidence of a longstanding large population size (Wegmann and Excoffier 2010) and is thought to be the oldest subspecies. The Eastern chimpanzee (*Pan troglodytes schweinfurthii*) emerged from the Central chimpanzees ancestors by a founder event and is the most recent sub-speciation event amongst chimpanzees (Hey 2010). Moving westward in Africa from the Central-Eastern chimpanzee distribution, two related subspecies are recognized: Nigerian-Cameroon chimpanzee (*Pan troglodytes ellioti*) and Western chimpanzee (Gonder, et al. 2011). The four chimpanzee subspecies show enough genetic distance for speculation that they may be in the early stages of speciation (Hey 2010). Additionally, bonobos (*Pan paniscus*) are a sister species to the common chimpanzee in the *Pan* genus, and reside in the Congo Basin between the current Eastern and Central chimpanzee distributions.

Similar to humans, copy number variation (CNV) of amplicon genes has been observed in chimpanzees and bonobos using florescence *in situ* hybridization (FISH) experiments (Repping, et al. 2006; Schaller, et al. 2010). However, these experiments focused on copy number of *CDY* and *DAZ*. Furthermore, a recent analysis of primate X-chromosome revealed that amplicons enriched for testis-specific genes are the target of hard-selective sweeps, suggesting that male-fertility traits may be the targets of strong selective forces in chimpanzees (Nam, et al. 2015). Despite these recent advancements, there are limited data of Y-linked amplicon content in the *Pan* lineage and given the dramatic structural changes identified in humans, we suspected the subspecies might harbor lineage specific amplicon content not present in the panTro4 reference.

A challenge in the completion of a mammalian genome sequence is an accurate assembly of the amplicon regions of the Y-chromosome. Amongst the great apes, all four genera (Gorilla, Pongo, Pan, and Homo) have high-resolution reference genome assemblies, however, only the chimpanzee and human have fully sequenced Y-chromosomes (Hughes, et al. 2010; Locke, et al. 2011; Scally, et al. 2012; Skaletsky, et al. 2003). The amplicon regions of Y-chromosomes have been assembled with single-haplotype iterative mapping and sequencing (SHIMS), a clone-by-clone approach that identifies overlapping inserts through sequence-tagged sites (STS) content assayed with PCR, sanger sequencing of BAC end sequences, and restriction enzyme patterns (Hughes and Page 2015). At this point, the extensive work required to assemble the amplicon sequence of Y-chromosomes limits our perspective of Y-chromosome diversity across great ape species and populations despite recent sequencing efforts (Prado-Martinez, et al. 2013). Alternatively, a draft gorilla Y chromosome has been mostly assembled through flow sorting Y-chromosomes and applying short-and long read genome and transcriptome sequencing (Tomaszkiewicz, et al. 2016). However, with this approach the ampliconic structure of the Y-chromosome remains poorly resolved.

To address this gap in our understanding of chimpanzee Y-chromosomes, we developed a method to estimate copy-number of amplicons and their nested genes from next-generation sequencing data. Our method utilizes short read data by counting a pre-defined set of amplicon k-mers (oligmers of length k) identified in the reference that match strict stringency thresholds for conservation within amplicon families and a lack of non-specificity. We first applied our method to an exploratory set of high coverage males from the 1000 Genomes (1KG) dataset (Genomes Project, et al. 2012). We validated computationally derived copy number estimates of amplicons with digital droplet PCR (ddPCR) and a found strong concordance in results. We next applied our method to a comprehensive analysis of a diverse set of 11 chimpanzee and bonobo male samples sequenced by the Great Ape Genome Project GAGP (Prado-Martinez, et al. 2013). We show that three chimpanzee subspecies evaluated here (Western Chimpanzee, Eastern Chimpanzee, and Nigerian-Cameroon Chimpanzee) and bonobos have distinctive Y-chromosome amplicon content, including variable copy-number of AZFb and AZFc orthologs, *RBMY* and *DAZ*.

## Material and Methods

### Sample selection and data processing

We analyzed 11 individuals previously sequenced by the Great Ape Genome Project with evidence of minimal sequence contamination (Prado-Martinez, et al. 2013). Selected samples included two bonobos (*Pan paniscus*: A919_Desmond, A925_Bono), and nine chimpanzees from three subspecies including four Nigerian-Cameroon chimpanzees (*Pan troglodytes ellioti*: Akwaya_Jean, Basho, Damian, Koto), two Eastern chimpanzees (*Pan troglodytes schweinfurthii*: 100037_Vincent, A910_Bwambale), two Western chimpanzees (*Pan troglodytes verus*: 9668_Bosco, Clint), and one Western/Central-hybrid (9730_Donald) (Table S2). Analysis utilized reads mapped to the panTro-2.1.4 (UCSC panTro4) assembly processed as previously described. We included human sample HGDP00222 (Pathan) (Martin, et al. 2014) as an out-group. Human reads were mapped to panTro4 using bwa (Li and Durbin 2009) (version 0.5.9) with options aln –q 15 –n 0.01 and sampe -o 1000 and processed as previously described using Picard (version 1.62) and the Genome Analysis Toolkit (version 1.2-65) (McKenna, et al. 2010). Our human data additionally includes publically available read alignments (merge of mapped and unmapped bam files) from 9 human individuals from the 1000 genomes project (NA12342, NA11994, NA12155, NA18623, NA18622, NA18636, NA19213, NA18519, NA19119). Reads from were aligned to human reference assembly hg19 and processed following 1KG consortium guidelines(Genomes Project, et al. 2012).

### Amplicon Analysis

The Y chromosome sequence assembly in panTro4 differs slightly from that originally reported (Hughes, et al. 2010). We therefore determined the boundaries of amplicon and palindrome units defined by Hughes et al. using the dotplot software Gepard (Krumsiek, et al. 2007). We assessed copy-number of each amplicon family using Quick-mer, a novel pipeline for paralog-specific copy-number analysis (Shen et al, in preparation, https://github.com/KiddLab/QuicK-mer). Briefly, QuicK-mer utilizes the Jellyfish-2 (Marcais and Kingsford 2011) program to efficiently tabulate the depth of coverage of a predefined set of k-mer sequences (here, k=30) in a set of sequencing reads. The resulting k-mer counts are normalized to account for effects of local GC percentage on read depth (Alkan, et al. 2009). For this analysis, we utilized two sets of k-mers: 30 mers determined to be unique throughout the genome, and k-mers determined to be specific to each individual amplicon family. For example, since there are four copies of the “red” amplicon in panTro4, we identified 93,200 k-mers present in each copy of the red amplicon and otherwise absent from the reference.

To identify candidate k-mers specific to each amplicon family, the sequences of the amplicon family were extracted from the reference sequence and then blatted to confirm that the sequence was found at the expected copy number in the reference. Next, 30bp k-mers found across the amplicon set were tabulated using Jellyfish-2 and those k-mers present at the expected copy-number were extracted. Genome wide unique k-mer candidates were identified based on Jellyfish analysis of the genome reference assembly. In all cases, Jellyfish-2 was ran in such a way that a k-mer and its reverse complement were considered to be identical. We applied a series of three quality control filters to remove k-mers that share identity with other sequence in the reference genome. First, we predefined a mask consisting of 15mers that are overrepresented in the genome. Any candidate 30mer intersecting with this mask was eliminated. Second, all matches against the reference genome within 2 substitutions were identified using mrsFAST (Hach, et al. 2010) and any kmer with >100 matches was eliminated. Finally, all mapping locations within an edit distance of 2, including indels, were identified using mrFAST (Alkan, et al. 2009; Xin, et al. 2013) and any kmer with >100 matches was filtered from the analysis. For *RBMY*, we were unable to identify k-mers at the expected copy number in the reference due to prevalent pseudogenization. We increased the expected copy number in the panTro4 reference to 10 from 6 to maximize k-mer coverage.

Depth for genome-wide unique and amplicon-specific kmers was then determined using Quickmer (Shen and Kidd 2015). GC normalization curves were calculated based on autosomal regions not previously identified as copy-number variable. K-mer depths were converted to copy-number estimates by dividing by the average depth of Y chromosome k-mers that passed the callable genome mask filter in the X-degenerate region on the chimpanzee Y. Prior to analysis all Y chromosome k-mers with a depth >0 in the female chimpanzee sample Julie_A959 (SAMN01920535) were removed. K-mer by position plots and heirachical clustering was created with R.

### ddPCR Validation of Quick-mer

CNV calls for select amplicons were validated in genomic DNA from human samples NA12342, NA11994, NA12155, NA18623, NA18622, NA18636, NA19213, NA18519, NA19119 (Coriell, Camden, NJ) using Droplet Digital PCR technology (ddPCR) (BioRad, Hercules, CA). We digested 1 ug of DNA at 37C overnight in a 50ul reaction of 1xNEB4 with 2.5U of BsmI and HindIII. The following morning, an additional 2U of each enzyme was added and the digest continued for another hour. After incubation, the reaction was diluted with 50ul of H_2_O and 8ul was used for ddPCR. Primer/probe assays were designed with Primer3Plus software http://www.bioinformatics.nl/cgi-bin/primer3plus/primer3plus.cgi using settings recommended for ddPCR by BioRad and synthesized at manufacturer's recommended primer/probe ratio with a 6-FAM (Fluorescein) label on the target probes, a HEX label on the reference probes, and an Iowa Black Quencher on both probes (Integrated DNA Technologies, Coralville, Iowa). Multiple primer/probe sets were designed for each locus and reference then the optimal set selected based on droplet cluster formation with minimal rain and agreement with locus CNV in a reference genome (Supplemental Table 1). For ddPCR workflow, a 20ul mixture containing 0.9 uM primer, 0.25 uM probe, 8 ul (80 ng) of digested DNA, and 1x Bio-Rad ddPCR Supermix for Probes (no dUTP) (BioRad, Hercules, CA) was emulsified, droplets were transferred to 96 well reaction plate, and plate was heat-sealed with foil using manufacturer's recommended reagents and conditions. We amplified the samples with the following PCR cycling protocol: 10 minutes at 95°C, 40 cycles of 30 seconds at 94°C and a 1 minute extension at 60°C, followed by 10 minutes at 98°C and a hold at 8°C. After PCR, droplets were read on droplet reader and data was analyzed using manufacturer's recommended reagents. 'Positive' droplets were identified by fluorescence intensity and thresholds were determined manually for each experiment.

### Phylogenetic Analysis

To identify a robust set of variants, we imposed a series of regional and site-level filters similar to the procedure utilized for the human MSY (Poznik, et al. 2013). First, we generated a regional callability mask, which defines the regions across the chromosomes that yield reliable genotype calls with short read sequence data. We calculated average filtered read depth and the MQ0 ratio (the number of reads with a mapping quality of zero divided by total read depth) in contiguous 1kb windows based on the output of the GATK Unified Genotyper ran in emit all sites mode. We computed an exponentially-weighted moving average (EWMA) across the windows and removed regions that deviated from a narrow envelope that excluded the tails of the read depth and MQ0 distributions.

We next applied a series of site-level filters to the individual positions that passed the regional masking (Supplemental Figure 2). We excluded positions with MQ0 fraction < 0.10, that contained missing genotypes in at least one sample, or contained a site where the maximum likelihood genotype was heterozygous in at least one sample. Of the remaining sequence, we plotted the distribution of site-level depths and removed positions with depths at the tail ends of the distribution. We excluded all triallelelic sites and further filtered variants within 5bp of indels. Separate regional callability masks and site-level filters were generated for the human and the combined chimp-bonobo samples and positions that failed either mask were dropped from the analysis. The combined call set consisted of 4,233,540 callable basepairs on the Y-chromosome. A neighbor-joining tree (500 bootstraps) of the MSY callable sites was generated in MEGA 6.06 using the Tamura-Nei substitution model (Tamura, et al. 2013).

## Results

### Validation in Human Reference Populations

We first applied our method to five AZFb and AZFc amplicons, noted for recurrent deletions in azoospermia cases. We mapped the positions of the yellow, red, blue, teal, and green amplicons from a Y chromosome (hg19) self-alignment represented as a dot-plot. Stretches of self identify (~150-500kb) within the AZFc region were annotated as amplicons (Supplemental Table 2A). In total, we identified two copies of the yellow amplicon, four copies of the red amplicon, four copies of the blue amplicon, two copies of the teal amplicon, and three copies of the green amplicon. Our amplicon definitions are nearly identical to those previously reported by Hughes et al., though one minor difference is that we mapped amplicon boundaries where homology is shared across all members of the family (i.e. all members within a family are close to equal in amplicon length and the copy number of an amplicon should be consistent by position based on the hg19 assembly). We extracted sequence within our amplicon boundaries and generated a set of k-mers conserved within and specific to each amplicon family, gene, and spacer (see methods). This predefined reference set of k-mers was used as input for the paralog sensitive k-mer counting pipeline, Quick-mer (Shen and Kidd 2015). Quick-mer counts the depth of a set of k-mers from short-read data and estimates the copy number relative to a specified reference sequence of a known copy number. As our copy-number reference sequence, we used the X-degenerate region found on the Y-chromosome, where massive CNV is infrequent and we expected to be at a copy-number of one on most Y-haplotypes.

We applied Quick-mer and our pre-defined set of amplicon k-mers to a diverse set of high coverage samples (>6x) from the 1KG datasets. Our 1KG set includes nine samples representing 3 clades of the Y-chromosome tree: E, R, and O. The E, R, and O Y chromosomes are commonly found amongst European, African, and Asian peoples, respectively. While DNA or short read data from the biological reference sample (RP-11) is not publically available, data reported in the literature confirms that most individuals carry the same copy-number as the reference AZFc haplotype (Repping, et al. 2006).

Applying Quick-mer to the study population revealed estimates matching the reference AZFc structure in 9 of 9 samples (Supplemental Figure 1). To validate our Quick-mer estimates, we used digital droplet PCR (ddPCR) to replicate our results. We designed our ddPCR primers within blocks of contiguous amplicon k-mers and used X-degenerate sequence as a copy number one control. After excluding yellow amplicon ddPCR results due to excessive “rain”, we were able to validate Quick-mer results in the remaining four amplicon families (Table 1).

**Table 1:**
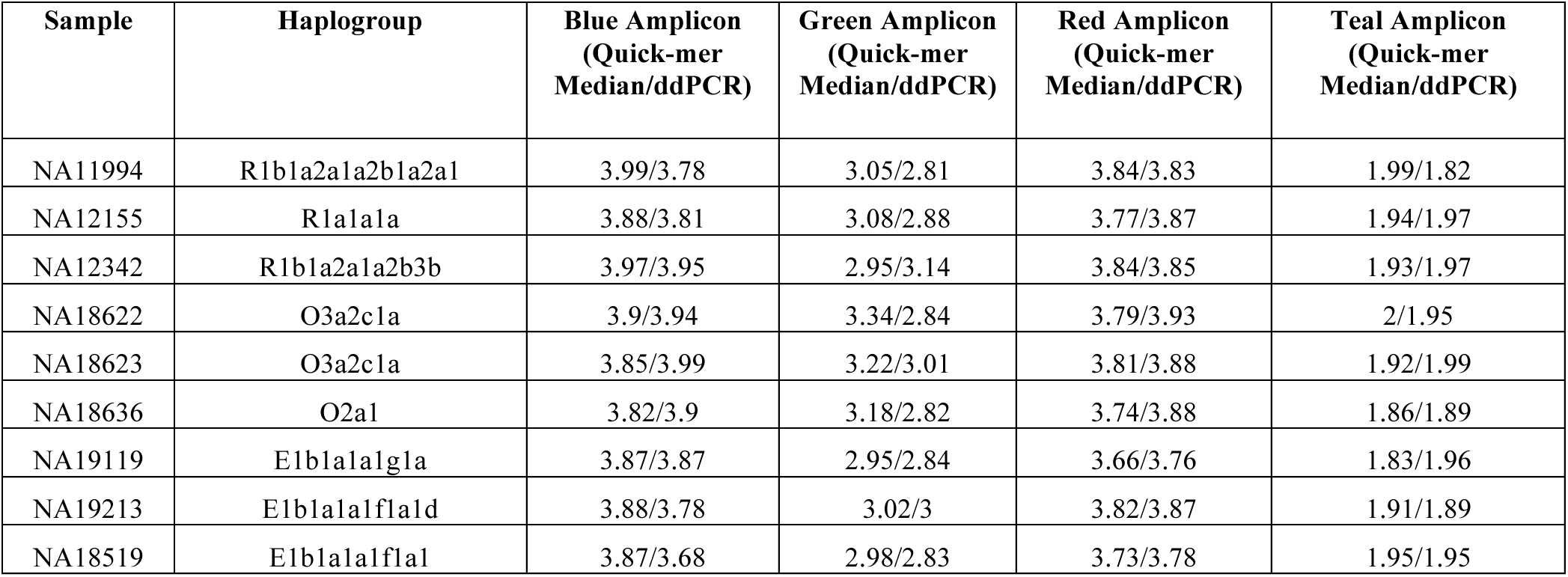
ddPCR Validation of Quick-mer AZFc Copy-Number Estimates in 1KG samples.

### Pan Amplicon Diversity

We applied the same procedure outlined above for mapping amplicons on the panTro4 version of the Y-chromosome. In total, we identified the 51 units across 10 amplicon families previously annotated by Hughes et al. (Supplemental Table 2B). Additionally, we mapped the unique “spacer” sequence between palindrome arms and ampliconic-gene positions (based on refSeq and ensembl) from the UCSC genome browser (Karolchik, et al. 2003) (Supplemental Table 2C). As a control, we compared Quickmer amplicon copy number estimates estimated from Clint to the panTro4 reference (Table 2A and Table 2b). After rounding median k-mer values to the nearest integer, we find strong agreement between the values, suggesting that Quick-mer provides accurate estimates of amplicon copy number. However, a few disagreements between our estimate of Clint's copy number and the reference sequence are evident. Quick-mer estimates Clint’s copy number of “violet arms” based on the median depth to be 11.54 (S.D. = 2.53) compared with the 13 represented in the reference assembly. Our copy number estimates are also different from the reference in the *CDY* (Clint: 3.98 vs. panTro4: 5), *VCY* (Clint: 0.77 vs. panTro4: 2), and *RBMY* (Clint 6 vs. panTro4: 10.91) genes. These differences likely arise from truncated copies of the genes and noise in our analysis due to the relatively short length of these genes. However, assembly errors in the panTro4 reference may also be a contributing factor. We also measured copy number a human sample (HGDP00222) based on the same 30-mers used to estimate chimpanzee copy-number. Many of our copy number estimates of HGDP00222 match the structures present in the human reference sequence as reported (Hughes, et al. 2010). Observed differences between the copy-number present in the human assembly (hg19) and HGDP00222 reflect a loss of chimpanzee-human homology for some 30-mers or amplicon polymorphism differences between HGDP00222 and the hg19 reference.

**Table 2.** Copy-number of amplicons and ampliconic genes by Pan population A. Amplicons, B. Ampliconic genes.

|  | Hughes et al. 2010<br>(Human) | Hughes et al. 2010<br>(Chimpanzee) | Clint | Human | Eastern<br>Average* | Western<br>Average* | Nigerian-Cameroon<br>Average* | Bonobo<br>Average* |
| --- | --- | --- | --- | --- | --- | --- | --- | --- |
| aqua arms | 0 | 6 | 5.49 | 0 | 5.9 | 5.53 | 13.43 | 10.16 |
| aqua spacer | 0 | 3 | 3.22 | 0 | 2.91 | 3.11 | 6.93 | 6.3 |
| blue arms | 4 | 4 | 3.84 | 2.66 | 1.98 | 3.85 | 3.86 | 2.2 |
| green arms | 3 | 2 | 1.87 | 3.43 | 1.04 | 1.91 | 1.93 | 1.72 |
| orange arms | 0 | 3 | 2.7 | 0 | 3.16 | 2.77 | 3.73 | 1.05 |
| other |  |  | 0.93 | 0 | 0.93 | 0.92 | 0.92 | 0.94 |
| pink arms | 0 | 4 | 3.86 | 0.95 | 5.99 | 3.81 | 11.87 | 13.47 |
| purple arms<sup>+</sup> | 0 | 3 | 2.82 | 0 | 0 | 2.78 | 4.74 | 0 |
| red arms | 4 | 4 | 3.92 | 4.39 | 2.01 | 3.89 | 3.92 | 2.26 |
| red spacer | 2 | 2 | 1.87 | 1.52 | 1.06 | 1.92 | 1.96 | 1.09 |
| teal arms | 2 | 7 | 6.68 | 0.95 | 9.66 | 5.46 | 12.4 | 11.13 |
| teal spacer | 0 | 3 | 2.84 | 0 | 4.89 | 2.21 | 5.71 | 4.76 |
| violet arms | 0 | 13 | 11.54 | 1.14 | 20.08 | 10.63 | 28.57 | 26.44 |
| violet spacer | 0 | 7 | 6.84 | 0 | 10.29 | 6.22 | 14.72 | 13.52 |
| yellow arms | 2 | 5 | 4.87 | 2.65 | 5.02 | 4.9 | 5.08 | 3.34 |
| yellow spacer | 0 | 2 | 1.83 | 2.3 | 2.09 | 1.95 | 1.95 | 1.04 |
| yellow spacer | 0 | 3 | 2.69 | 2.72 | 3.1 | 2.72 | 3.15 | 2.1 |

\*Average Median Value, <sup>+</sup> Additional purple amplicon mapped in the panTro4 assembly

*B. Ampliconic genes*
|  | **Hughes et al. 2010 (Human)** | **Hughes et al. 2010 (Chimpanzee)** | **Clint** | **Human** | **Eastern (Average)** | **Western (Average)** | **Nigerian-Cameroon (Average)** | **Bonobo (Average)** |
| --- | --- | --- | --- | --- | --- | --- | --- | --- |
| *BPY2* | 3 | 2 | 1.81 | 3.61 | 1.1 | 1.86 | 2.07 | 1.79 |
| *CDY* | 4 | 5 | 3.98 | 4 | 5.86 | 4.51 | 5.41 | 3.59 |
| *DAZ* | 4 | 4 | 4.37 | 0.19 | 1.76 | 4.11 | 3.85 | 2.26 |
| *RBMY* | 6 | 6 | 10.91 | 10.32 | 14.34 | 9.16 | 24.12 | 32.28 |
| *TSPY* | 35 | 6 | 5.46 | 38.85 | 16.6 | 10.27 | 26.35 | 32.1 |
| *VCY* | 2 | 2 | 0.77 | 0 | 0.44 | 1.01 | 1.41 | 0.05 |

UCSC Genome Browser Gene Identifier: *CDY* (NM\_001145039), *DAZ* (ENSPTRG00000022523), *BPY2* (NM\_004678.2), *RBMY* (NM\_001006121.2), *TSPY* (NM\_001077697.2), *VCY* (NM\_004679.2)

To confirm the Y-chromosomes of our chimpanzee samples are consistent with the autosomal inferred phylogeny, we constructed a SNP callset from the panTro4 Y chromosome. Following the procedure reported by Poznik et al. (Poznik, et al. 2013), we used alignment quality scores to identify regions of the Y chromosome amendable to variant calling. We identified 4.2Mb of sequence in the X-degenerate regions for variant calling with chimpanzee and human samples. A neighbor-joining tree confirms that the 3 chimpanzee subspecies form monophyletic clades (Figure 1). The Western Chimpanzee Donald from the GAGP was previously reported to have Nigerian-Cameroon Chimpanzee admixture. However, from our phylogenetic tree we are able to verify his Y chromosome is of Western Chimpanzee origin and therefore we included him in our analysis as a Western Chimpanzee. We also observe a deep divergence between the bonobo samples, nearly equal to the common ancestor of the chimpanzee in age. As point of comparison, our tree closely resembles a recent MSY inferred chimpanzee phylogeny (Hallast, et al. 2016).

**Fig. 1:**
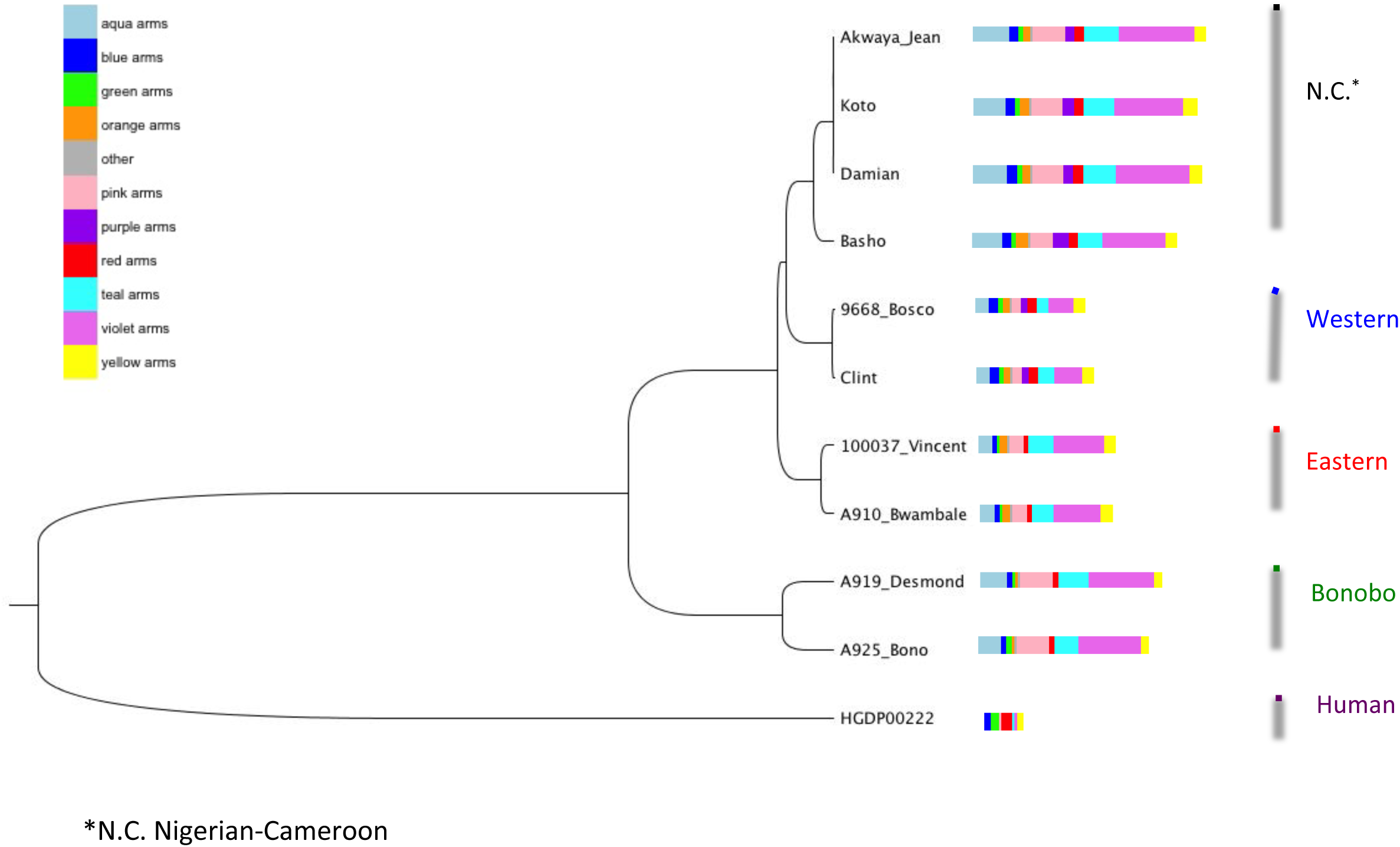
Y Chromosome Tree and Amplicon Copy Number. The phylogenetic relationship among sequenced *Pan* and human Y chromosomes is illustrated with neighbor joining tree using a Tamura-Nei substitution model. The size of the color bars to the right of the tree indicate relative median copy number of each amplicons estimated per sample. Note: the order of the color bars does not reflect the spatial distribution of amplicons.

We see substantial variance in the copy number of amplicons, spacers, and ampliconic-genes across the chimpanzee subspecies and bonobos (Table 2A, Table 2B, Supplemental Table 3A, Supplemental 3B). However, within individual *Pan* lineages we observe an overall conservation of copy number in the ampliconic regions (Figure 1). The teal and violet palindromes found within the 2.2Mb palindrome array near the centromere are highly variable in copy number(Hughes, et al. 2010). This correlated with an increase of the *RBMY* gene across samples (median k-mer depth; Western: 9.16, Eastern: 14.34, Nigerian-Cameroon: 24.12, bonobos: 32.28) (Supplemental Figure 3J and Supplemental Figure 3K). We note that our *RBMY* estimates are likely inflated due to psuedogenization of the gene on the chimpanzee Y-chromosome, as evident in the panTro4 assembly (see methods). The Western chimpanzees are also comparably lacking in pink arms and the adjacent *TSPY* gene: the median copy number of the pink amplicon (3.81) and the *TSPY* gene (10.27) is one-fourth and one-half of the estimates in Nigerian-Cameroon chimpanzees (pink amplicon: 11.87, *TSPY*: 26.35) and bonobos (pink amplicon: 13.47, *TSPY*: 32.10), respectively (Supplemental Figure 3T). However, copy number *TSPY* increases by position in all samples, therefore it is likely our estimates of *TSPY* copy number (based on k-mer counts) may also include truncated of copies of the gene.

We find that *DAZ* and the red amplicon ranges in copy number between two (Eastern and bonobos) and four (Nigerian-Cameroon and Westerns) copies. Interestingly, within the red amplicon we observe two exon-spanning deletions in the human and bonobo samples that share a common break point located in intron 15 of *DAZ* (Figure 2A, Supplemental Figure 3Q). A similar event is observed in the purple arms (Figure 2D), where there is amplification of copy number in the Eastern chimpanzees and reduction in the bonobos on the 5' of a breakpoint, and the reverse of the pattern on the 3' end.

**Fig. 2:**
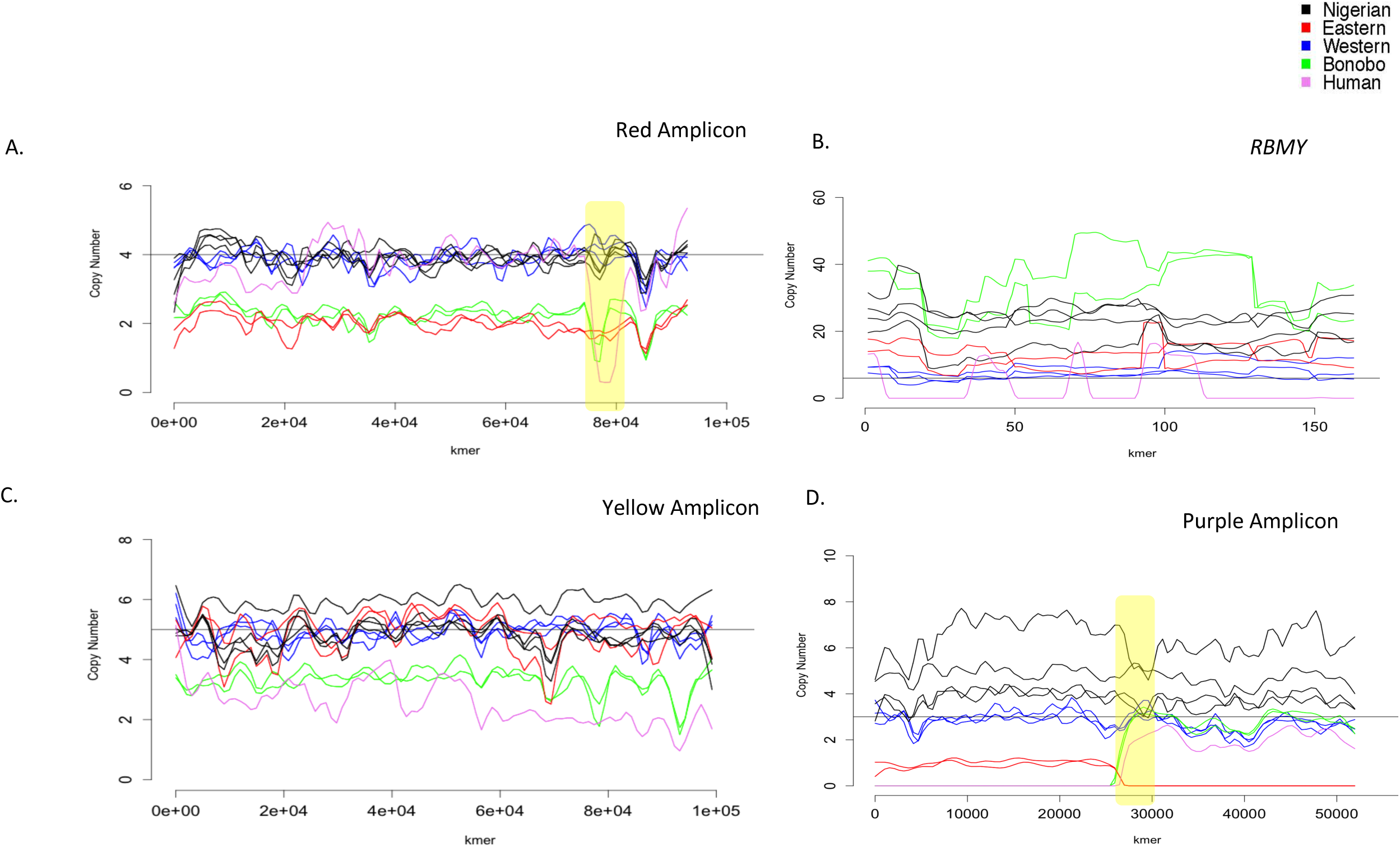
Chimpanzee and Bonobo Amplicon Copy Number by K-mer. Amplicon copy number are plotted against k-mer position. Each line represents a single sample colored by species/subspecies and smoothed with lowess function. Copy number of the PanTro4 reference is drawn with a solid black line. Breakpoints of interest mentioned in the text are boxed in yellow. *A. Red Amplicon B. RBMY C. Yellow Amplicon D. Purple Amplicon*.

## Discussion

Our data reveal that amplicons continue to evolve in chimpanzees, leading to distinct gene content found in the Eastern, Western, and Nigerian-Cameroon chimpanzee Y-chromosomes. This finding represents an important step in our understanding of the evolution of sex chromosomes, specifically in the early stages of primate speciation. As a point of comparison, there are no subspecies classifications for bonobos and here we find limited amplicon copy number diversity between individuals, consistent with a previous report (Schaller, et al. 2010). This is a surprising observation in light of the high level of nucleotide diversity we detect on the bonobo Y-chromosome (Figure 1) and may suggest that amplicon CNV may be the result of selection of male-fertility in the chimpanzee lineages.

The *DAZ* locus is a candidate fertility factor and thought to be at least partially responsible for the azoosperima phenotype in AZFc deletions. Molecular clock analyses of the *DAZ* locus have revealed considerable evolution since its transposition to the primate Y chromosome from chromosome 3 38.5 mya (Saxena, et al. 1996). Subsequently, the Y chromosome paralog duplicated 33 million years ago, prior to the Old World Monkey - Ape split, and today it is found in two copies in the rhesus macaque (Hughes, et al. 2012). Comparisons between the chimpanzee and human reference sequence revealed four copies in both assemblies. However, comparative analysis of the bordering palindrome sequences reveals that these are the result of independent amplifications in chimpanzee and human (Hughes, et al. 2010; Skaletsky, et al. 2003). As noted in a previous study, copy number of *DAZ* is polymorphic amongst the common chimpanzees (Schaller, et al. 2010). Here we find that Eastern chimpanzees and bonobos each have two copies of *DAZ* and the red amplicon, while Western and Nigerian-Cameroon have four copies. With respect to copy number, we hypothesize that the bonobos and Eastern represent the ancestral state of the *DAZ* locus in the Pan lineage. Our results support a scenario where the second *DAZ* duplication event evident in the panTro4 assembly is the result of a duplication that occurred early in the formation of the Western/Nigerian-Cameroon clade. Furthermore, the *DAZ* RNA-recognition motif shows substantial positive selection in the chimpanzee reference assembly relative to the human and rhesus macaque assemblies. However, if the amplification of *DAZ* is unique to the Western/Nigerian-Cameroon lineages, we suspect that Y-chromosome selection may be higher in these clades. Hierarchical clustering of the chimpanzee and bonobo samples (based on fold change of amplicon copy number) reveals a Western/Nigerian-Cameroon chimpanzee and bonobo/eastern Chimpanzee divergence that supports this hypothesis (Supplemental Figure 4). Interestingly, this tree structure persists even after the removal of the red amplicon from the analysis as this result is also driven by the blue and purple amplicons (Table 2A). However, we are limited in power to infer phylogeny from our copy-number data due to low counts of unique amplicon structures (n=10) and samples (n=11). Additionally, such analyses are complicated by the potential for recurrent copy-number changes along these lineages. If male fertility has influenced the diversification of the common chimpanzee, the *DAZ* duplication we observe in the Western and Nigerian-Cameroon clade may have played a critical role. Further studies are required to determine if the *DAZ* is under positive selection in the Central/Eastern chimpanzees and bonobos.

Additionally, copy number of *RBMY* displayed conspicuous variability across our samples (Table 2B). While copy number of the *RBMY* k-mers in the common chimpanzees is relatively constant by position, we observe abrupt changes in bonobo copy number at switch points within the gene (Figure 2B). In chimpanzees, multiple copies of *RBMY* are nested within the “palindrome array” of violet and teal amplicons, which display amplification in chimpanzee populations and bonobos relative to Western chimpanzees (Table 3A). These results suggest that the palindrome array is subject to structural rearrangement on the *Pan* Y-chromosome. Furthermore, the conservation of *RBMY* on the Y-chromosome since marsupials (divergence from placental mammals was ~160 mya (Luo, et al. 2011) suggests that this locus has an important male-specific function (Delbridge, et al. 1997). This stands in contrast to *DAZ*, which moved from the autosomes to the Y-chromosome exclusively in the primate lineage (Gromoll, et al. 1999). Additionally, the testis exhibit extensive alternative splicing of *RBMY* and it is one of the few genes identified to activate testis-specific splicing events (Liu, et al. 2009; Yeo, et al. 2004). The *RBMY* CNVs may result in variable expression of testis-specific isoforms across *Pan* and could contribute to population specific male-fertility traits. We also characterized unique Western chimpanzee sequence within the non-repetitive sequence in the ampliconic region termed “other” (Supplemental Figure 7E). We speculate that there may be subspecies-specific amplicons not assayed here. Future studies of the chimpanzee Y-chromosomes would benefit from high-quality MSY assemblies specific for subspecies to identify patterns of gain and loss of unique sequence.

We found that our k-mer counting approach is optimal for detecting CNV of amplicons and more limited with small genes and highly repetitive arrays such as the *TSPY*. Our method of assaying CNVs of ampliconic genes is also limited in that it does not distinguish between intact and pseudogenized copies. Furthermore, divergence between the reference and the analyzed samples may reduce the accuracy of our approach. However, the bonobos represent a divergence from the panTro4 reference of approximately ~1-2 mya and we find that the k-mer by position plots show a minimal increase in noise (Prado-Martinez, et al. 2013). In contrast, the analyzed human sample, which represents a ~6 mya divergence, displays uneven copy number by position across amplicons and genes. While the relative quantity of annotated amplicons generated from short read data is reported here, assemblies would provide the spatial diversity of amplicons. With assemblies in hand, one could recreate the non-homologous recombination events that gave rise to the structural diversity within *Pan* and further clarify the contribution of ampliconic sequence to subspecies evolution.

## Acknowledgments

This work was supported by the National Institutes of Health (grant number 1DP5OD009154). We thank Brenna Henn and Jacob Mueller for thoughtful comments on manuscript drafts.

## Supplemental Figure Legends

*Supplemental Fig. 1: Human Amplicon Copy Number by K-mer*

Plots of lowess smoothed copy number of ampliconic features by diagnostic k-mer are drawn on the left and histograms of copy number are drawn on the right. Sample population is indicated by line color and the solid black line represents expected copy number based on Hughes et al. 2010.

*Supplemental Fig. 2: Callability Mask for Identifying the X-degenerate Sites*

Exponentially weighted moving averages (EWMA) of read depth (blue line) and the mq0/unfiltered depth ratio (pink line) are plotted against position on the Y chromosome (PanTro4). Dashed lines represent maximum and minimum thresholds for filtered depth (green) and a maximum threshold for the mq0 ratio (red). Colored bars below plot indicate regions masked by the depth filter (blue), masked by the mq0 ratio filter (pink), excluded from the analysis (grey), and included (black) for further filtering steps.

*Supplemental Fig. 3: Chimpanzee and Bonobo Amplicon Copy Number by K-mer (All Amplicons)*

Plots of lowess smoothed copy number of ampliconic features by diagnostic k-mer are drawn on the left and histograms of copy number are drawn on the right. Samples are colored by population/species and the solid black line represents expected copy number based on Hughes et al. 2010.

*Supplemental Fig. 4: Hierarchical Clustering*

Dendrogram of *Pan* samples computed based on relative distance of copy number depth of ampliconic arms. Distance based on copy number between two samples *i* and *j* across *n* ampliconic features was calculated by the following formula:

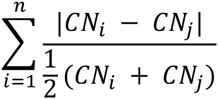

A. Amplicon arms only (spacers and genes excluded) B. Amplicon arms only (without the red arms)

## References

Alkan C, Kidd JM, Marques-Bonet T, Aksay G, Antonacci F, Hormozdiari F, Kitzman JO, Baker C, Malig M, Mutlu O, Sahinalp SC, Gibbs RA, Eichler EE 2009. Personalized copy number and segmental duplication maps using next-generation sequencing. Nat Genet 41: 1061–1067.

Bachtrog D 2013. Y-chromosome evolution: emerging insights into processes of Y-chromosome degeneration. Nat Rev Genet 14: 113–124. doi: 10.1038/nrg3366

Delbridge ML, Harry JL, Toder R, O'Neill RJ, Ma K, Chandley AC, Graves JA 1997. A human candidate spermatogenesis gene, RBM1, is conserved and amplified on the marsupial Y chromosome. Nat Genet 15: 131–136. doi: 10.1038/ng0297-131

Genomes Project C, Abecasis GR, Auton A, Brooks LD, DePristo MA, Durbin RM, Handsaker RE, Kang HM, Marth GT, McVean GA 2012. An integrated map of genetic variation from 1,092 human genomes. Nature 491: 56–65. doi: 10.1038/nature11632

Gonder MK, Locatelli S, Ghobrial L, Mitchell MW, Kujawski JT, Lankester FJ, Stewart CB, Tishkoff SA 2011. Evidence from Cameroon reveals differences in the genetic structure and histories of chimpanzee populations. Proc Natl Acad Sci U S A 108: 4766–4771. doi: 10.1073/pnas.1015422108

Gromoll J, Weinbauer GF, Skaletsky H, Schlatt S, Rocchietti-March M, Page DC, Nieschlag E 1999. The Old World monkey DAZ (Deleted in AZoospermia) gene yields insights into the evolution of the DAZ gene cluster on the human Y chromosome. Hum Mol Genet 8: 2017–2024.

Gu W, Zhang F, Lupski JR 2008. Mechanisms for human genomic rearrangements. Pathogenetics 1: 4. doi: 10.1186/1755-8417-1-4

Hach F, Hormozdiari F, Alkan C, Hormozdiari F, Birol I, Eichler EE, Sahinalp SC 2010. mrsFAST: a cache-oblivious algorithm for short-read mapping. Nat Methods 7: 576–577. doi: 10.1038/nmeth0810-576

Hallast P, Maisano Delser P, Batini C, Zadik D, Rocchi M, Schempp W, Tyler-Smith C, Jobling MA 2016. Great ape Y Chromosome and mitochondrial DNA phylogenies reflect subspecies structure and patterns of mating and dispersal. Genome Res 26: 427–439. doi: 10.1101/gr.198754.115

Hey J 2010. The divergence of chimpanzee species and subspecies as revealed in multipopulation isolation-with-migration analyses. Mol Biol Evol 27: 921–933. doi: 10.1093/molbev/msp298

Hughes JF, Page DC 2015. The Biology and Evolution of Mammalian Y Chromosomes. Annu Rev Genet 49: 507–527. doi: 10.1146/annurev-genet-112414-055311

Hughes JF, Skaletsky H, Page DC 2012. Sequencing of rhesus macaque Y chromosome clarifies origins and evolution of the DAZ (Deleted in AZoospermia) genes. Bioessays 34: 1035–1044. doi: 10.1002/bies.201200066

Hughes JF, Skaletsky H, Pyntikova T, Graves TA, van Daalen SK, Minx PJ, Fulton RS, McGrath SD, Locke DP, Friedman C, Trask BJ, Mardis ER, Warren WC, Repping S, Rozen S, Wilson RK, Page DC 2010. Chimpanzee and human Y chromosomes are remarkably divergent in structure and gene content. Nature 463: 536–539. doi: 10.1038/nature08700

Karolchik D, Baertsch R, Diekhans M, Furey TS, Hinrichs A, Lu YT, Roskin KM, Schwartz M, Sugnet CW, Thomas DJ, Weber RJ, Haussler D, Kent WJ, University of California Santa C 2003. The UCSC Genome Browser Database. Nucleic Acids Res 31: 51–54.

Kidd JM, Graves T, Newman TL, Fulton R, Hayden HS, Malig M, Kallicki J, Kaul R, Wilson RK, Eichler EE 2010. A human genome structural variation sequencing resource reveals insights into mutational mechanisms. Cell 143: 837–847. doi: 10.1016/j.cell.2010.10.027

Krumsiek J, Arnold R, Rattei T 2007. Gepard: a rapid and sensitive tool for creating dotplots on genome scale. Bioinformatics 23: 1026–1028. doi: 10.1093/bioinformatics/btm039

Lahn BT, Pearson NM, Jegalian K 2001. The human Y chromosome, in the light of evolution. Nat Rev Genet 2: 207–216. doi: 10.1038/35056058

Li H, Durbin R 2009. Fast and accurate short read alignment with Burrows-Wheeler transform. Bioinformatics 25: 1754–1760. doi: btp324 [pii] 10.1093/bioinformatics/btp324

Liu Y, Bourgeois CF, Pang S, Kudla M, Dreumont N, Kister L, Sun YH, Stevenin J, Elliott DJ 2009. The germ cell nuclear proteins hnRNP G-T and RBMY activate a testis-specific exon. PLoS Genet 5: e1000707. doi: 10.1371/journal.pgen.1000707

Locke DP, Hillier LW, Warren WC, Worley KC, Nazareth LV, Muzny DM, Yang SP, Wang Z, Chinwalla AT, Minx P, Mitreva M, Cook L, Delehaunty KD, Fronick C, Schmidt H, Fulton LA, Fulton RS, Nelson JO, Magrini V, Pohl C, Graves TA, Markovic C, Cree A, Dinh HH, Hume J, Kovar CL, Fowler GR, Lunter G, Meader S, Heger A, Ponting CP, Marques-Bonet T, Alkan C, Chen L, Cheng Z, Kidd JM, Eichler EE, White S, Searle S, Vilella AJ, Chen Y, Flicek P, Ma J, Raney B, Suh B, Burhans R, Herrero J, Haussler D, Faria R, Fernando O, Darre F, Farre D, Gazave E, Oliva M, Navarro A, Roberto R, Capozzi O, Archidiacono N, Della Valle G, Purgato S, Rocchi M, Konkel MK, Walker JA, Ullmer B, Batzer MA, Smit AF, Hubley R, Casola C, Schrider DR, Hahn MW, Quesada V, Puente XS, Ordonez GR, Lopez-Otin C, Vinar T, Brejova B, Ratan A, Harris RS, Miller W, Kosiol C, Lawson HA, Taliwal V, Martins AL, Siepel A, Roychoudhury A, Ma X, Degenhardt J, Bustamante CD, Gutenkunst RN, Mailund T, Dutheil JY, Hobolth A, Schierup MH, Ryder OA, Yoshinaga Y, de Jong PJ, Weinstock GM, Rogers J, Mardis ER, Gibbs RA, Wilson RK 2011. Comparative and demographic analysis of orang-utan genomes. Nature 469: 529–533. doi: 10.1038/nature09687

Longepied G, Saut N, Aknin-Seifer I, Levy R, Frances AM, Metzler-Guillemain C, Guichaoua MR, Mitchell MJ 2010. Complete deletion of the AZFb interval from the Y chromosome in an oligozoospermic man. Hum Reprod 25: 2655–2663. doi: 10.1093/humrep/deq209

Luo ZX, Yuan CX, Meng QJ, Ji Q 2011. A Jurassic eutherian mammal and divergence of marsupials and placentals. Nature 476: 442–445. doi: 10.1038/nature10291

Marcais G, Kingsford C 2011. A fast, lock-free approach for efficient parallel counting of occurrences of k-mers. Bioinformatics 27: 764–770. doi: 10.1093/bioinformatics/btr011

Martin AR, Costa HA, Lappalainen T, Henn BM, Kidd JM, Yee MC, Grubert F, Cann HM, Snyder M, Montgomery SB, Bustamante CD 2014. Transcriptome sequencing from diverse human populations reveals differentiated regulatory architecture. PLoS Genet 10: e1004549. doi: 10.1371/journal.pgen.1004549 PGENETICS-D-13-02851 [pii]

McKenna A, Hanna M, Banks E, Sivachenko A, Cibulskis K, Kernytsky A, Garimella K, Altshuler D, Gabriel S, Daly M, DePristo MA 2010. The Genome Analysis Toolkit: a MapReduce framework for analyzing next-generation DNA sequencing data. Genome Res 20: 1297–1303. doi: gr.107524.110 [pii] 10.1101/gr.107524.110

Nam K, Munch K, Hobolth A, Dutheil JY, Veeramah KR, Woerner AE, Hammer MF, Great Ape Genome Diversity P, Mailund T, Schierup MH 2015. Extreme selective sweeps independently targeted the X chromosomes of the great apes. Proc Natl Acad Sci U S A 112: 6413–6418. doi: 10.1073/pnas.1419306112

Poznik GD, Henn BM, Yee MC, Sliwerska E, Euskirchen GM, Lin AA, Snyder M, Quintana-Murci L, Kidd JM, Underhill PA, Bustamante CD 2013. Sequencing Y chromosomes resolves discrepancy in time to common ancestor of males versus females. Science 341: 562–565. doi: 341/6145/562 [pii] 10.1126/science.1237619

Prado-Martinez J, Sudmant PH, Kidd JM, Li H, Kelley JL, Lorente-Galdos B, Veeramah KR, Woerner AE, O'Connor TD, Santpere G, Cagan A, Theunert C, Casals F, Laayouni H, Munch K, Hobolth A, Halager AE, Malig M, Hernandez-Rodriguez J, Hernando-Herraez I, Prufer K, Pybus M, Johnstone L, Lachmann M, Alkan C, Twigg D, Petit N, Baker C, Hormozdiari F, Fernandez-Callejo M, Dabad M, Wilson ML, Stevison L, Camprubi C, Carvalho T, Ruiz-Herrera A, Vives L, Mele M, Abello T, Kondova I, Bontrop RE, Pusey A, Lankester F, Kiyang JA, Bergl RA, Lonsdorf E, Myers S, Ventura M, Gagneux P, Comas D, Siegismund H, Blanc J, Agueda-Calpena L, Gut M, Fulton L, Tishkoff SA, Mullikin JC, Wilson RK, Gut IG, Gonder MK, Ryder OA, Hahn BH, Navarro A, Akey JM, Bertranpetit J, Reich D, Mailund T, Schierup MH, Hvilsom C, Andres AM, Wall JD, Bustamante CD, Hammer MF, Eichler EE, Marques-Bonet T 2013. Great ape genetic diversity and population history. Nature 499: 471–475. doi: nature12228 [pii] 10.1038/nature12228

Repping S, van Daalen SK, Brown LG, Korver CM, Lange J, Marszalek JD, Pyntikova T, van der Veen F, Skaletsky H, Page DC, Rozen S 2006. High mutation rates have driven extensive structural polymorphism among human Y chromosomes. Nat Genet 38: 463–467. doi: 10.1038/ng1754

Rozen S, Skaletsky H, Marszalek JD, Minx PJ, Cordum HS, Waterston RH, Wilson RK, Page DC 2003. Abundant gene conversion between arms of palindromes in human and ape Y chromosomes. Nature 423: 873–876. doi: 10.1038/nature01723

Rozen SG, Marszalek JD, Irenze K, Skaletsky H, Brown LG, Oates RD, Silber SJ, Ardlie K, Page DC 2012. AZFc deletions and spermatogenic failure: a population-based survey of 20,000 Y chromosomes. Am J Hum Genet 91: 890–896. doi: 10.1016/j.ajhg.2012.09.003

Saxena R, Brown LG, Hawkins T, Alagappan RK, Skaletsky H, Reeve MP, Reijo R, Rozen S, Dinulos MB, Disteche CM, Page DC 1996. The DAZ gene cluster on the human Y chromosome arose from an autosomal gene that was transposed, repeatedly amplified and pruned. Nat Genet 14: 292–299. doi: 10.1038/ng1196-292

Scally A, Dutheil JY, Hillier LW, Jordan GE, Goodhead I, Herrero J, Hobolth A, Lappalainen T, Mailund T, Marques-Bonet T, McCarthy S, Montgomery SH, Schwalie PC, Tang YA, Ward MC, Xue Y, Yngvadottir B, Alkan C, Andersen LN, Ayub Q, Ball EV, Beal K, Bradley BJ, Chen Y, Clee CM, Fitzgerald S, Graves TA, Gu Y, Heath P, Heger A, Karakoc E, Kolb-Kokocinski A, Laird GK, Lunter G, Meader S, Mort M, Mullikin JC, Munch K, O'Connor TD, Phillips AD, Prado-Martinez J, Rogers AS, Sajjadian S, Schmidt D, Shaw K, Simpson JT, Stenson PD, Turner DJ, Vigilant L, Vilella AJ, Whitener W, Zhu B, Cooper DN, de Jong P, Dermitzakis ET, Eichler EE, Flicek P, Goldman N, Mundy NI, Ning Z, Odom DT, Ponting CP, Quail MA, Ryder OA, Searle SM, Warren WC, Wilson RK, Schierup MH, Rogers J, Tyler-Smith C, Durbin R 2012. Insights into hominid evolution from the gorilla genome sequence. Nature 483: 169–175. doi: 10.1038/nature10842

Schaller F, Fernandes AM, Hodler C, Munch C, Pasantes JJ, Rietschel W, Schempp W 2010. Y chromosomal variation tracks the evolution of mating systems in chimpanzee and bonobo. PLoS One 5. doi: 10.1371/journal.pone.0012482

Shen F, Kidd J 2015. QuicK-mer: A rapid paralog sensitive CNV detection pipeline. bioRxiv. doi: 10.1101/028225

Skaletsky H, Kuroda-Kawaguchi T, Minx PJ, Cordum HS, Hillier L, Brown LG, Repping S, Pyntikova T, Ali J, Bieri T, Chinwalla A, Delehaunty A, Delehaunty K, Du H, Fewell G, Fulton L, Fulton R, Graves T, Hou SF, Latrielle P, Leonard S, Mardis E, Maupin R, McPherson J, Miner T, Nash W, Nguyen C, Ozersky P, Pepin K, Rock S, Rohlfing T, Scott K, Schultz B, Strong C, Tin-Wollam A, Yang SP, Waterston RH, Wilson RK, Rozen S, Page DC 2003. The male-specific region of the human Y chromosome is a mosaic of discrete sequence classes. Nature 423: 825–837. doi: 10.1038/nature01722

Tamura K, Stecher G, Peterson D, Filipski A, Kumar S 2013. MEGA6: Molecular Evolutionary Genetics Analysis version 6.0. Mol Biol Evol 30: 2725–2729. doi: 10.1093/molbev/mst197

Tomaszkiewicz M, Rangavittal S, Cechova M, Sanchez RC, Fescemyer HW, Harris R, Ye D, O'Brien PC, Chikhi R, Ryder OA, Ferguson-Smith MA, Medvedev P, Makova KD 2016. A time- and cost-effective strategy to sequence mammalian Y Chromosomes: an application to the de novo assembly of gorilla Y. Genome Res 26: 530–540. doi: 10.1101/gr.199448.115

Wegmann D, Excoffier L 2010. Bayesian inference of the demographic history of chimpanzees. Mol Biol Evol 27: 1425–1435. doi: 10.1093/molbev/msq028

Xin H, Lee D, Hormozdiari F, Yedkar S, Mutlu O, Alkan C 2013. Accelerating read mapping with FastHASH. BMC Genomics 14 Suppl 1: S13. doi: 10.1186/1471-2164-14-S1-S13

Yeo G, Holste D, Kreiman G, Burge CB 2004. Variation in alternative splicing across human tissues. Genome Biol 5: R74. doi: 10.1186/gb-2004-5-10-r74

